# Plasticity in mosquito size and thermal tolerance across a latitudinal climate gradient

**DOI:** 10.1101/2024.06.05.597660

**Authors:** Kelsey Lyberger, Johannah E. Farner, Lisa Couper, Erin A. Mordecai

## Abstract

Variation in heat tolerance among populations can determine whether a species is able to cope with ongoing climate change. Such variation may be especially important for ectotherms whose body temperatures, and consequently, physiological processes, are regulated by external conditions. Additionally, differences in body size are often associated with latitudinal clines, thought to be driven by climate gradients. While studies have begun to explore variation in body size and heat tolerance within species, our understanding of these patterns across large spatial scales, particularly regarding the roles of plasticity and genetic differences, remains incomplete. Here, we examine body size, as measured by wing length, and thermal tolerance, as measured by the time to immobilization at high temperatures (“thermal knockdown”), in populations of the mosquito *Aedes sierrensis* collected from across a large latitudinal climate gradient spanning 1300 km (34-44 °N). We find that mosquitoes collected from lower latitudes and warmer climates were more tolerant of high temperatures than those collected from higher latitudes and colder climates. Moreover, body size increased with latitude and decreased with temperature, a pattern consistent with James’ rule, which appears to be a result of plasticity rather than genetic variation. Our results suggest that warmer environments produce smaller and more thermally tolerant populations.

## Introduction

Understanding how species vary across different environments is a fundamental goal in ecology, with implications for biodiversity, ecosystem functioning, and the responses of species to climate change. This is particularly relevant for ectotherms, such as mosquitoes, whose physiological processes are directly influenced by external environmental conditions. Consistent patterns of variation in traits such as body size across latitudinal gradients have been established across species, where larger organisms are associated with colder climates at high latitudes or elevations (Bergmann’s rule, Blackburn et al. 1999). Equally important is establishing the extent to which this pattern also applies within species (i.e., James’ rule, James 1970). Both patterns may result from a combination of evolution (genetic variation between and within species) and phenotypic plasticity (variation among individuals that is in response to environmental conditions). Understanding these patterns is important as body size is linked to many physiological and ecological functions including thermal regulation, resource use, and reproductive success. While these rules were initially established in endotherms, previous work has found they also apply to ectotherms in part as a result of warmer temperatures driving faster development rates and reducing oxygen availability (e.g., in Diptera; Baranov et al. 2022).

Temperature is a fundamental driver of the physiological processes of ectotherms. Numerous studies in insects have documented the temperature-size rule, where organisms reared at higher temperatures often develop smaller body sizes (Atkinson 1994; Davidowitz et al. 2004; Garrad et al. 2016), although there are exceptions such as in cicadas (Koyama et al. 2015; Nguyen et al. 2020). This phenomenon is particularly relevant in the context of climate change, as rising temperatures could lead to shifts in body size distributions and affect survival, reproduction, and behavior that governs species interactions such as feeding patterns (Angilletta 2009; Klockmann et al. 2017).

In addition to body size per se, thermal tolerance, which may depend on body size, is another critical trait that determines a species resilience to climate change. Decades of research has demonstrated that, *between* terrestrial ectotherm species, heat tolerance decreases minimally with latitude (Deutsch et al. 2008; Sunday et al. 2019). However, only a few studies have investigated thermal tolerance across latitude *within* a species (Klepsatel et al. 2013; Overgaard et al. 2014; Ruybal et al. 2016). These studies have typically used laboratory-reared populations rather than individuals collected in the wild. The degree to which a natural climate gradient leads to variation in body size and thermal tolerance within a species remains an open question.

This study aims to address this gap by examining the intraspecific variation in body size and thermal tolerance of the western tree hole mosquito, *Aedes sierrensis,* across a latitudinal climate gradient. This species represents an ideal system as its range spans western North America. By analyzing both individuals that were collected as pupae in the field and those that were reared entirely in the laboratory from populations spanning a 1,300 km range from 34 to 44 °N, this research seeks to answer: (1) How much intraspecific variation in size exists across a latitudinal cline and how correlated is it with temperature? (2) Are patterns in size, if observed, driven by environmental or genetic differences? (3) Do populations from warmer, lower latitudes have higher thermal tolerance, and is this affected by individual body size? By answering these questions, we aim to identify how plasticity in body size and thermal tolerance traits respond to environmental variation and enhance our understanding of how ectotherms might respond to changing climatic conditions.

## Materials and methods

### Field collection and study system

*Aedes sierrensis* inhabit water-filled treeholes across the Pacific Coast of North America. In this region, eggs hatch following fall and winter rains and mosquitoes develop into adults before the treeholes dry out in summer. Eggs diapause during the dry season and, at high latitudes, 4th instar larvae diapause during the wet season (Jordan and Bradshaw 1978). This species is thought to be univoltine across its range, which is ideal as species with changing voltinism are expected to produce a sawtooth pattern rather than smooth cline of increasing size with latitude (Kivelä et al. 2011).

*Aedes sierrensis* were sampled from populations ranging from southern California to northern Oregon, covering 1,300 km and the majority of this species’ range (Figure 1). For the main plasticity experiment, samples from 13 populations were collected between January and April of 2023, with later sampling dates at higher latitudes. We only used individuals collected in the pupal stage to ensure they had completed feeding as larvae and reached full potential adult size (as individuals do not feed at the pupal life stage). Once collected, pupae were brought to the lab and allowed to develop into adults at 21°C and a 13 h:11 h light:dark cycle. We then conducted ‘thermal knockdowns’ to assay thermal tolerance and measured wing length as a measure of body size, both of which are described further below. To determine the extent to which body size is driven by plasticity rather than genetic differences, we also present an analysis of data from an experiment conducted in 2022, in which larvae from populations spanning a similar latitudinal gradient were collected and reared for a generation under laboratory conditions across a range of temperatures (see methods in Couper et al. 2024). Here, we report body size for 9 populations reared at 4 temperatures including 13, 17, 24, and 28 (°C).

**Figure 1.**
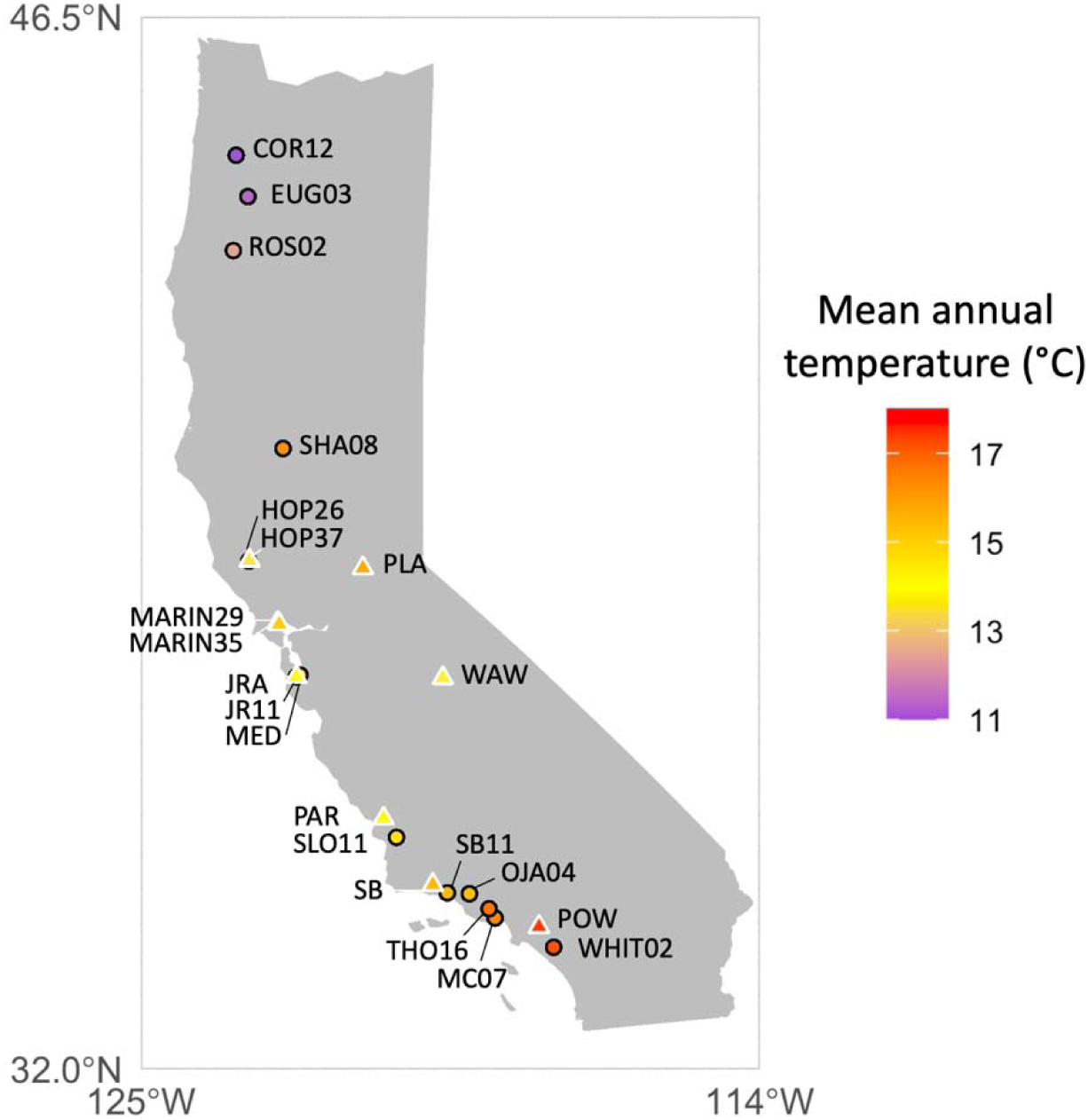
Map of the locations in California and western Oregon where mosquito pupae were collected for the plasticity study (black circles) and where mosquito larvae were collected for the common garden study (white triangles). Points are colored by the mean annual temperature.

We excluded one population (EUG) from the original study, which produced no adults at 13°C, and two temperature treatments: 5°C, in which larvae failed to develop into pupae, and 32°C, in which the majority of larvae and pupae died before emerging as adults. For this species, depending on the life-history trait, the estimated thermal minima range from 0 to 8°C, thermal optima range from 12 to 27°C, and thermal maxima range from 30 to 35°C (Couper et al. 2024). We measured the wing length of all individuals and the thorax length of individuals from the population MAR35 as further described in the methods below. Coordinates and sample sizes are given in Table 1.

**Table 1.** Population site names and localities, sample sizes (N) of *Aedes sierrensis* males and females, and climate information on mean annual temperature.

| Population | Location | Latitude | Longitude | Males | Females | T<sub>annual</sub> (°C) |
| --- | --- | --- | --- | --- | --- | --- |
| *Samples in the main thermal knockdown plasticity experiment* |  |  |  |  |  |  |
| WHIT02 | Trabuco Canyon, CA | 33.68171 | -117.66436 | 20 | 13 | 17.4 |
| MC07 | Malibu Creek, CA | 34.08479 | -118.70553 | 20 | 20 | 16.5 |
| THO16 | Thousand Oaks, CA | 34.21223 | -118.81246 | 10 | 14 | 16.8 |
| OJA04 | Ojai, CA | 34.41842 | -119.16085 | 20 | 20 | 15.4 |
| SB11 | Santa Barbara, CA | 34.43218 | -119.55338 | 20 | 20 | 15.5 |
| SLO11 | Lopez Lake, CA | 35.19735 | -120.45880 | 8 | 14 | 14.6 |
| JR11 | Jasper Ridge, CA | 37.41106 | -122.23690 | 20 | 20 | 14.0 |
| MED01 | Stanford, CA | 37.43076 | -122.17537 | 20 | 20 | 14.8 |
| HOP26 | Hopland, CA | 39.00157 | -123.08195 | 20 | 20 | 14.4 |
| SHA08 | Shasta, CA | 40.55275 | -122.47546 | 20 | 20 | 16.3 |
| ROS02 | Roseburg, OR | 43.28008 | -123.35902 | 9 | 9 | 12.5 |
| EUG21 | Eugene, OR | 44.02215 | -123.09983 | 20 | 15 | 11.5 |
| COR12 | Corvallis, OR | 44.59196 | -123.30870 | 16 | 20 | 11.1 |
| *Samples reared at a common temperature* |  |  |  |  |  |  |
| POW | La Habra, CA | 33.9638888 | -117.92432 | 19 | 28 | 17.6 |
| SB | Santa Barbara, CA | 34.54276 | -119.81219 | 20 | 30 | 15.5 |
| PAR | Atascadero, CA | 35.455543 | -120.68839 | 16 | 28 | 14.1 |
| WAW | Yosemite Forks, CA | 37.391291 | -119.6331 | 16 | 27 | 13.7 |
| JRA | Jasper Ridge, CA | 37.40805 | -122.23703 | 16 | 17 | 13.9 |
| MARIN35 | Novato, CA | 38.1304913 | -122.54399 | 23 | 40 | 15.1 |
| MARIN29 | Novato, CA | 38.1525295 | -122.57082 | 15 | 14 | 15.0 |
| PLA | Auburn, CA | 38.9039747 | -121.05436 | 13 | 18 | 15.8 |
| HOP37 | Hopland, CA | 39.0099819 | -123.07222 | 25 | 20 | 13.6 |

### Thermal knockdown trials

Between 48-72 hours after emergence, we subjected adults to a thermal knockdown trial. This time was chosen so that males and females were likely to have reached sexual maturity. Although it is unknown when this occurs for *Aedes sierrensis*, in congeneric species males tend to reach maturity one to two days after emergence (Miyagi & Toma 1982, Chevone et al. 1976, Provost et al. 1961) and females can take a blood meal two days after eclosure (Lyimo & Takken 1993). Adults were starved prior to the trial as we found that sugar-feeding impacted individual thermal tolerance in preliminary trials.

To perform the thermal knockdown trial, we followed a modified version of standard methods for this assay developed for *Drosophila* (Jørgensen et al. 2019). Briefly, individual mosquitoes were housed in a sealed 5ml plastic vial and submerged in a temperature-controlled water bath. The mosquitoes were allowed to acclimate for 15 minutes at 28°C. Then, at 2-minute intervals, the temperature was raised by 0.5°C up to a maximum of 38°C. To account for any variation in this ramping procedure, we recorded the time 38°C was reached. Mosquitoes were watched continuously until they were motionless, at which time they were jostled to encourage movement. Once a mosquito was motionless for 5 seconds, the exposure time until knockdown was recorded. A maximum of 11 individuals were run per trial. All trials were conducted by the same observer to ensure consistency. After the trial, individuals were frozen at -80°C to be measured at a later time.

### Wing measurements

To assess mosquito body size, we measured individual wing length – a commonly used body size measure (Lyimo & Takken 1993; Ameneshewa and Service 1996; Blackmore and Lord 2000; Davis et al. 2016). To measure wing length, we dissected the right wing, and laid it on a microscope slide with a 2 mm micrometer to standardize size measurements. Because relative wing length has been known to vary between sexes and under different conditions in other insects (Azevedo et al. 1998, Rohner and Moczek 2020), we also measured thorax length on a subset of individuals to determine whether the wing to thorax ratio differed among sexes, populations, and temperature treatments. We measured the thorax of nearly all field-collected mosquitoes from all populations and lab-reared mosquitoes from the population MARIN35, excluding those which were damaged from the removal of the wing. Thorax length was measured as the distance between the most anterior edge of the thorax to the most posterior end of the scutellum when viewed from the side (Yeap et al. 2013). Images of the wings and thorax were captured using an iPhone camera mounted to an Olympus BX51 microscope. We processed the images using ImageJ, measuring the length from the alular notch to the tip of the wing margin, excluding the wing scales (Abramoff et al. 2004).

### Climate data

To capture the environmental conditions at the locations where the populations were collected (i.e., the source environment), we obtained historical climate data on mean annual temperature from the WorldClim dataset, which calculates average bioclimatic variables from 1970-2000 at a 1 km resolution (Fick and Hijmans 2017). The mean annual temperature at our sampling sites ranged from 11.1 to 17.6°C (Table 1). We use these measurements of mean annual temperature when referring to ’source temperature’ below.

### Statistical analysis

We conducted multiple linear mixed effect models (LMMs), implemented using the *‘lme4’* package in R, to analyze the relationships between temperature at the site of collection (source temperature), sex, size, population, block (defined below) and knockdown time (heat tolerance) (Bates et al. 2015). For the main plasticity experiment we ran two models. In the first, body size was the dependent variable, source temperature, sex, and their interaction were included as fixed effects, and population was included as a random effect. In the second, knockdown time was the dependent variable, size, source temperature, sex, and the interaction between source temperature and sex were included as fixed effects, and both population and block were included as random effects, where block represents the set of individuals run during the same knockdown trial.

We also modeled the size of mosquitoes reared at the same temperatures under laboratory conditions as the dependent variable. This model included source temperature, sex, the interaction of source temperature and sex, and the temperature at which the mosquitoes were reared as fixed effects and included population as a random effect. For all models, assumptions, including normality of residuals and homoscedasticity, were checked using diagnostic plots. We reported statistical significance based on likelihood ratio tests and partial eta-squared using the *‘effectsize’* package in R (Ben-Shachar et al. 2020). Because latitude and temperature were strongly correlated with each other, we chose to focus on temperature as this is expected to have a more direct relationship with size and knockdown time, and because some of our populations came from altitudes of up to 650 m. We reported the effects of latitude in the supplement. We included both latitude and temperature in our SEM analysis below.

To better account for the complex causal relationships among variables, we also applied a structural equation modeling (SEM) approach—a linear modeling approach that enables testing of direct and indirect effects on pre-assumed causal relationships. To reflect the nested structure of our data, we employed a multilevel SEM, using population as the clustering variable. We tested the following causal pathways. First, higher latitudes have cooler climates. Second, source temperature reduces the average size of individuals in a population (between-population size).

Third, size at the individual level (within-population size) is affected by sex, with females being larger. Fourth, knockdown time is directly influenced by source temperature, with higher temperatures producing more thermally tolerant mosquitoes. Fifth, knockdown time is indirectly influenced by size at the individual level and size at the population level (i.e., the average size of individuals in a population), with smaller organisms being less thermally tolerant. Therefore, source temperature is expected to have two opposing effects, by directly increasing knockdown time and indirectly decreasing knockdown time by reducing body size. We fit the model and estimated parameters using the ’*lavaan*’ package in R (Rosseel 2012). To determine the significance of the paths within our model, we relied on z-tests that challenge the hypothesis of non-significant path coefficients. The fit of our proposed model to the data was assessed using a chi-square (χ²) test, which measures the divergence between the expected variance-covariance matrix as implied by the model and the actual observed matrix.

## Results

A total of 223 males and 225 females were collected as pupae from across western North America, underwent a thermal knockdown trial, and were measured for wing length (Table 1, Figure 1). The average wing length was 2.94 mm, ranging from 1.93 mm to 4.23 mm. The generalized linear models showed that there was a significant reduction in size with increasing temperature at the collection site (Figure 2A; Table 2). Individuals from the coldest site were on average 34% larger than those from the warmest site, where these two sites are separated by 1,310 km and differed by 6°C in mean annual temperatures. Similarly, the effect of latitude was significant, with each degree of latitude leading to an estimated 0.06 mm increase in wing length (Table S1). Females were significantly larger than males, as most *Aedes* mosquitoes are known to be sexually dimorphic in size (Zacarés et al. 2018). The males from one warm site THO16 (Thousand Oaks, CA) were outliers in being relatively large for their sex and source temperature. The site SHA08 (Shasta, CA) was an outlier in being located at a relatively high latitude but having a warm climate; the size of the mosquitoes from that site reflected the climate. Across all populations there was also a significant interaction with sex in that females showed a stronger body size response to temperature than males. We found no significant effect of sex or source temperature on wing to thorax ratio (Figure 2B, Table S2). This result supports the use of wing length as a proxy for body size.

**Figure 2.**
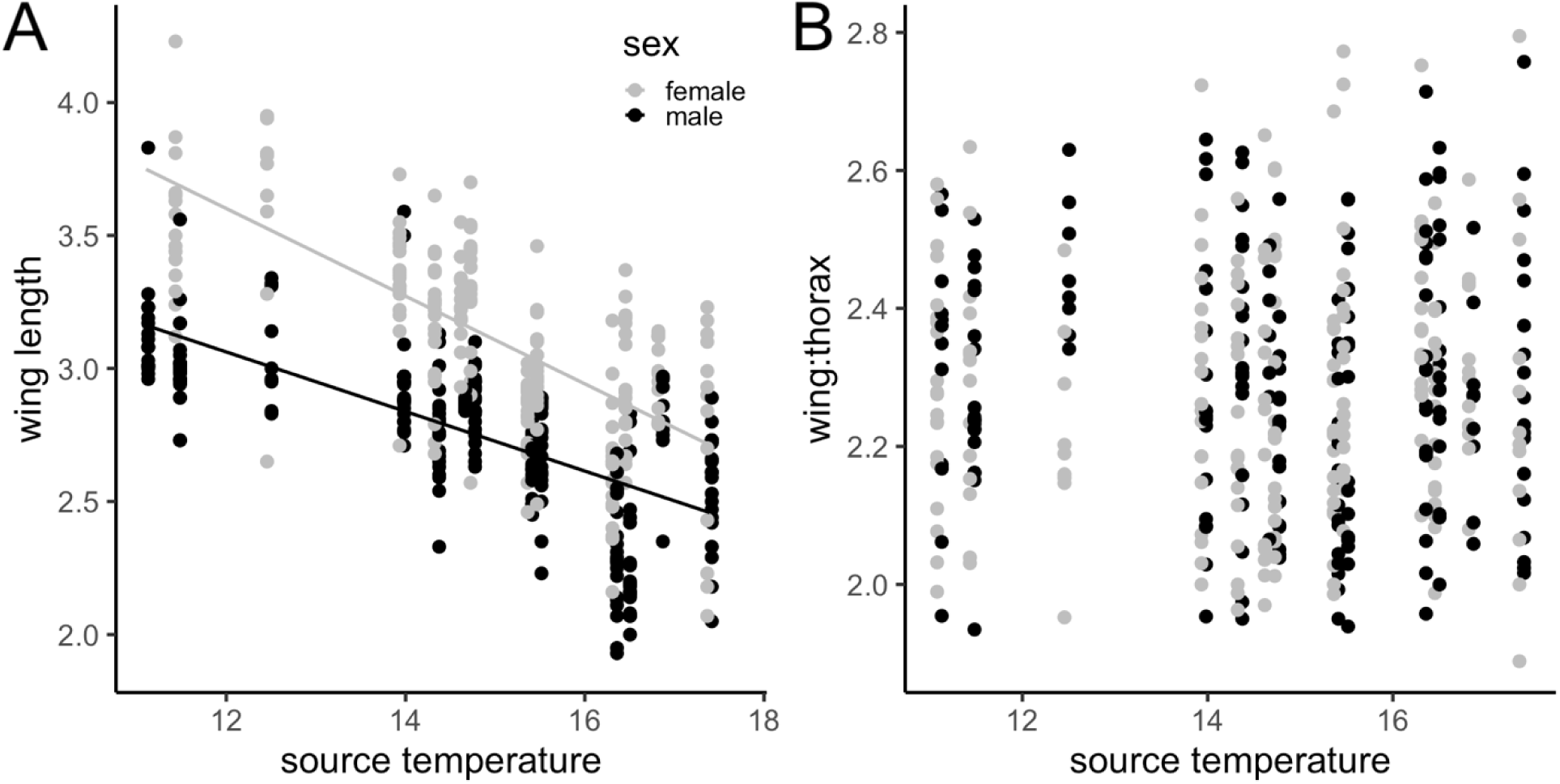
(A) For individuals collected directly from the field as pupae, body size decreased with the mean annual temperature at the collection site (estimate = -0.16, χ² = 21.7, P < 0.0001). (B) The wing to thorax ratio did not differ by sex (estimate = 0.01, χ² = 0.03, P = 0.86) or by collection site temperature (estimate = 0, χ² = 0.01, P = 0.91). Males are in black, and females are in gray.

**Table 2.** Results of the LMMs on the size and knockdown time of adults collected as pupae (N=448) and the size of adults reared their entire lives in controlled temperatures (N=385). Population was included as a random effect (sigma=0.13 for a, sigma=1.5 for b, sigma=0.08 for c). Block was included as a random effect (sigma=1.0 for b).

| Term | Estimate | SE | $\eta^2$ | df | $\chi^2$ | p-value |
| --- | --- | --- | --- | --- | --- | --- |
| (a) impact on size of field-collected individuals |  |  |  |  |  |  |
| temperature | -0.16 | 0.02 | 0.81 | 1 | 21.7 | <0.0001 |
| sex | -1.18 | 0.17 | 0.45 | 1 | 251.6 | <0.0001 |
| temperature x sex | 0.05 | 0.01 | 0.05 | 1 | 21.8 | <0.0001 |
| (b) impact on knockdown time |  |  |  |  |  |  |
| temperature | 1.14 | 0.29 | 0.37 | 1 | 8.2 | 0.0004 |
| sex | 6.68 | 2.83 | 0.003 | 1 | 1.5 | 0.22 |
| temperature x sex | 1.11 | 0.77 | 0.02 | 1 | 6.8 | 0.009 |
| size | -0.48 | 0.19 | 0.004 | 1 | 2.2 | 0.14 |
| (c) impact on size of lab-reared individuals |  |  |  |  |  |  |
| temperature (collected) | -0.03 | 0.02 | 0.18 | 1 | 1.8 | 0.18 |
| sex | -0.63 | 0.25 | 0.61 | 1 | 350.5 | <0.0001 |
| temperature (collected) x sex | -0.01 | 0.02 | 0.001 | 1 | 0.3 | 0.58 |
| temperature (reared) | -0.13 | 0.01 | 0.36 | 1 | 171.3 | <0.0001 |

In contrast, our laboratory study found that when reared at a common temperature in the lab (13, 17, 24, or 28°C), individuals showed no significant trend in wing length across populations, regardless of the temperature at the collection site (Figure 3A; Table 2). In that study, 205 females and 180 males were measured from a similar area across western North America (Table 1). We found that there was a significant difference in size between individuals reared at different temperatures, with higher temperatures leading to smaller size (Figure 3B; Table 2). Females were again significantly larger than males and there was no significant effect of sex or rearing temperature on wing to thorax ratio (Figure S1, Table S2). Summarizing across the two studies, body size was primarily driven by rearing temperature through the larval stages: mosquitoes collected after pupation in the field varied in size by source environment temperature (Figure 2), while those reared to pupation in the laboratory varied by rearing temperature but not by source environment temperature (Figure 3). This demonstrates that differences in size across populations are mostly driven by plastic effects of the environment rather than genetic differences.

**Figure 3.**
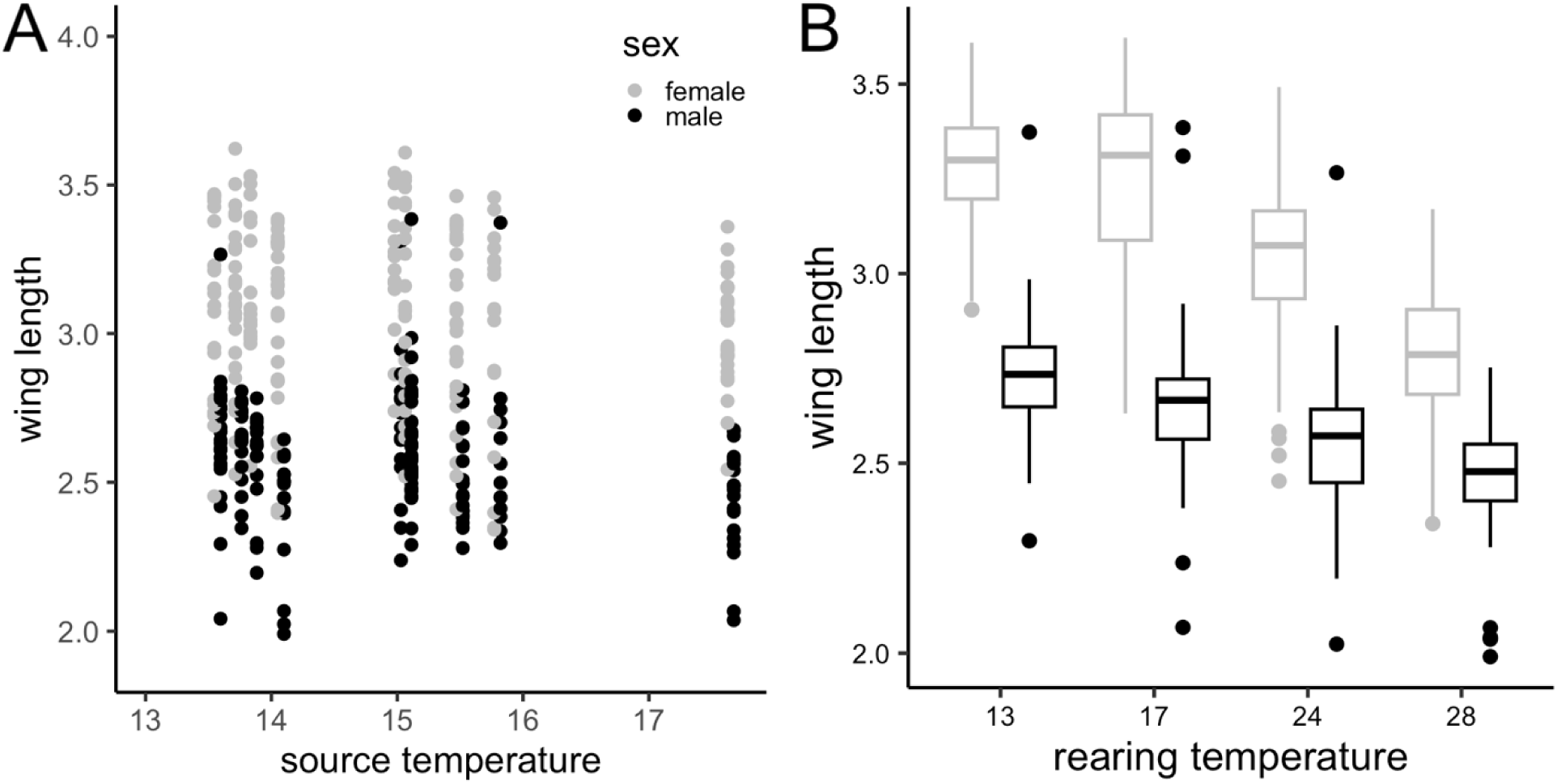
(A) For individuals raised to adulthood under laboratory conditions, the temperature at the collection site had no significant effect on body size (estimate = -0.03, χ² = 1.8, P = 0.18). (B In contrast, the temperature at which the mosquitoes were reared significantly impacted body size (estimate = -0.13, χ² = 171.3, P < 0.0001). Males are in black, and females are in gray.

We measured knockdown time as a measure of thermal tolerance for all individuals collected in the field as pupae, to assess the impact of source temperature and body size on thermal tolerance. Across all individuals and populations, the average time to knockdown was 43 minutes, ranging from 25 to 60 minutes. Knockdown time was significantly influenced by the temperature at the collection site, where each additional degree in mean annual temperature corresponded to an increase of more than a minute in knockdown time (Figure 4; Table 2).

**Figure 4.**
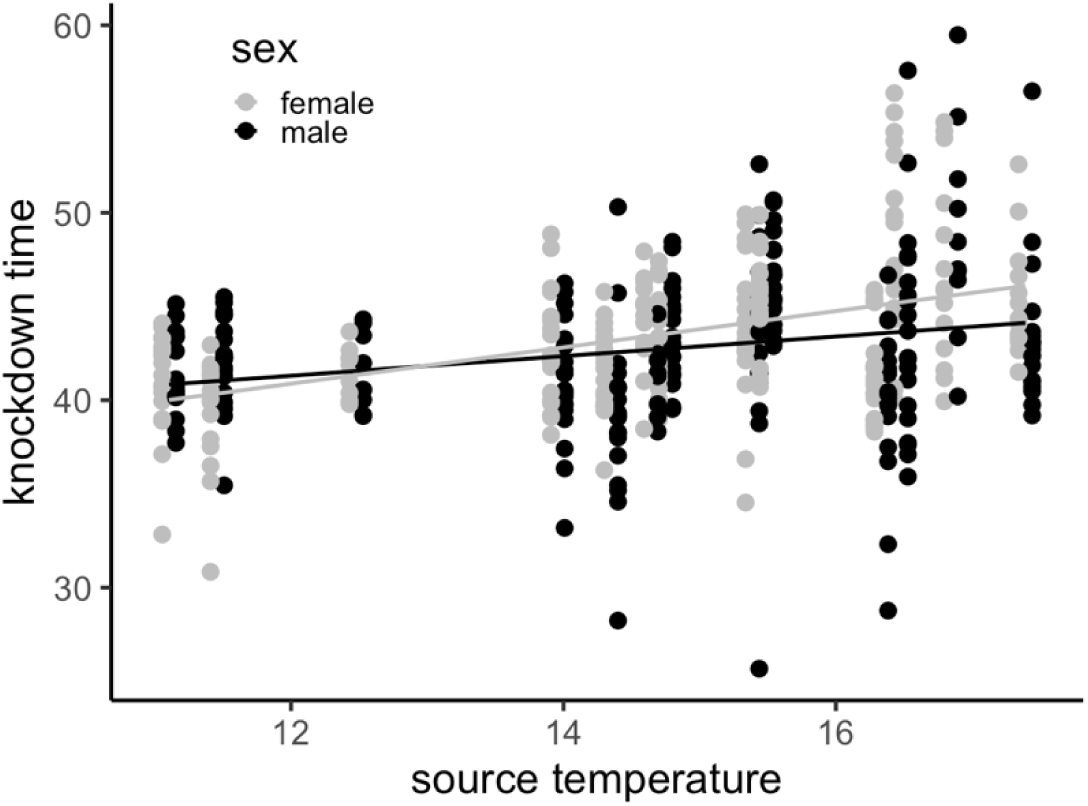
Knockdown time of mosquitoes collected from the field as pupae increased with mean annual temperature at the collection site (estimate = 1.14, χ² = 8.2, P = 0.0004). The strength of this relationship was stronger for females than males.

Individuals from the warmest site (‘WHIT02’) had on average a 7% longer knockdown time compared to individuals from the coldest site (‘COR12’). In the mixed effect model of knockdown time that included size, source temperature, sex, and the interaction between source temperature and sex, size was not a significant predictor (Table 2). There was a significant interaction between source temperature and sex, in which females from warmer sites exhibited longer knockdown times than males, whereas at cooler sites, females had shorter knockdown times than males (Table 2). Similar results were obtained when latitude is included in the model in place of temperature (Table S1).

Because we hypothesized that part of the effect of source temperature on knockdown time was driven by its effects on body size, we fit structural equation models that tested these hypothesized paths. This included pathways from latitude to source temperature to between-population size to knockdown time, from source temperature to knockdown time, and from sex to within-population size to knockdown time. All the hypothesized paths were significant (Figure 5). The chi-square goodness-of-fit test indicated that the hypothesized model provided a good fit to the data (χ²(3) = 5.226, P = 0.156). Latitude was negatively associated with temperature, and temperature, in turn, was negatively related to size. Sex was significantly associated with size, where males were smaller than females. Size was a significant predictor of knockdown time, with the between-population size having a larger effect than within-population size. Finally, the positive direct effect of source temperature on knockdown time was stronger than the negative indirect effect of source temperature through body size. Coefficients, standard errors, z-values, and p-values for each path are provided in Table 3.

**Figure 5.**
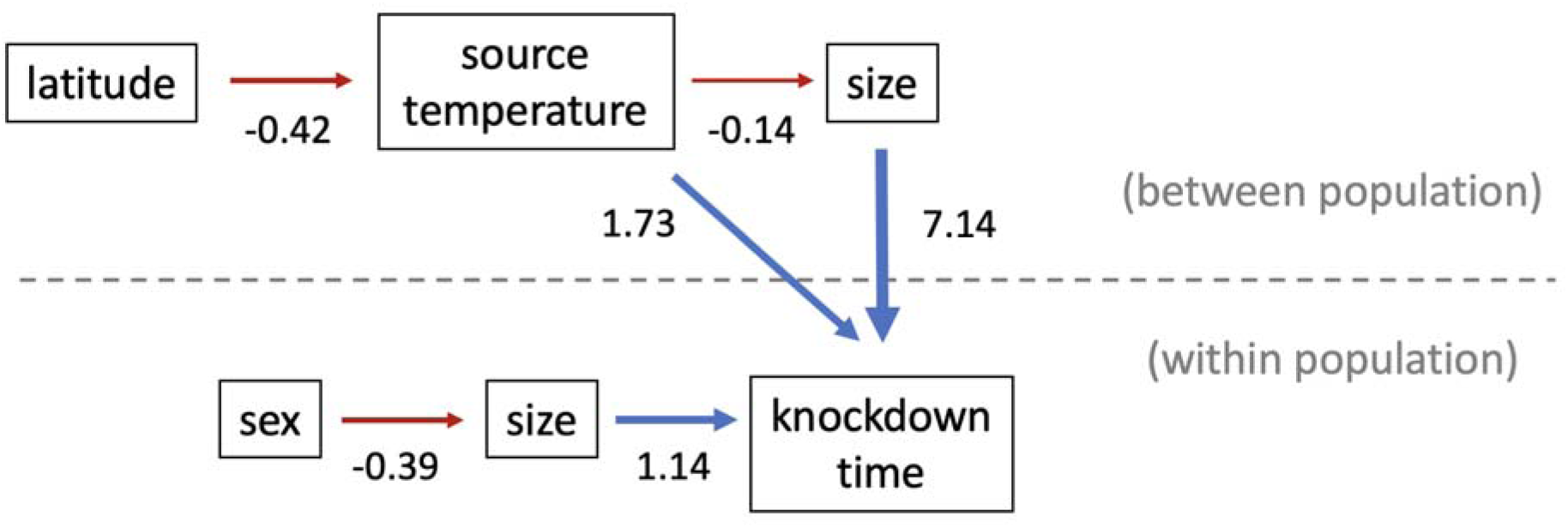
Source temperature has a direct positive effect on knockdown time and weaker indirect negative effect on knockdown time through a reduction in body size. The path coefficients of the multilevel structural equation model are shown next to the arrows. As the model uses population as a clustering variable, between population relationships (i.e., using a single average value for each population) are shown above the dashed line and within population relationships are below the dashed line. Red arrows denote negative relationships and blue arrows denote positive relationships.

**Table 3.** Parameter estimates of the structural equation model.

| Relationship | Estimate | SE | z-value | p-value |
| --- | --- | --- | --- | --- |
| *Between-population relationships* |  |  |  |  |
| temperature ~ latitude | -0.42 | 0.07 | -5.83 | 0.000 |
| size ~ temperature | -0.14 | 0.02 | -7.03 | 0.000 |
| knockdown ~ temperature | 1.73 | 0.48 | 3.58 | 0.000 |
| knockdown ~ size | 7.14 | 3.14 | 2.27 | 0.023 |
| *Within-population relationships* |  |  |  |  |
| size ~ sex | -0.39 | 0.02 | -18.19 | 0.000 |
| knockdown ~ size | 1.14 | 0.57 | 1.98 | 0.047 |

## Discussion

In this study, we found that mosquito body size (wing length) was smaller in populations from warmer source environments and lower latitudes, which aligns with the temperature – size rule and James’ rule as observed in other taxa (Figure 2A). This result is consistent with prior studies of arthropod size across latitudinal gradients, which are especially strong in aquatic species (Horne et al. 2015, 2017). Our observed differences in size were substantial and likely underlie important variation in populations’ life histories. For example, fecundity is positively associated with wing length in *Aedes sierrensis* (Washburn et al. 1989). Mosquitoes from the coldest climate were up to 34% larger than those from the warmest climate, which, if the relationship from Washburn et al. (1989) held, would translate into 46 more eggs laid or a 97% increase in clutch size.

By rearing a separate set of populations from a similar latitudinal and climatic gradient under common temperature conditions in the lab, we found evidence that these size differences are likely to be the result of plasticity across environments rather than genetic differences between populations. When populations were reared under laboratory conditions, the temperature at the collection site had no significant effect on size (Figure 3A) whereas the temperature at which they were reared in the lab had a strong negative effect on size (Figure 3B). This finding contrasts with several studies in *Drosophila*, which found evidence of rapid evolution of larger wings at higher latitudes decades after populations invaded latitudinal clines in western North America (Huey et al. 2000) and South America (Gilchrist et al. 2004). Although wing length is a commonly used proxy for body size in Diptera, wing and thorax length are just two of many possible measures. Body mass would provide a very direct metric; however, given the minimal mass of a mosquito, it was not feasible to measure body mass accurately.

In our study, when populations were collected as pupae (having completed all the feeding and growth that would contribute to adult body size in the field), temperature at the collection site was strongly correlated with body size, supporting existing evidence from laboratory studies that warmer temperatures accelerate development rates and reduce body size (Atkinson 1995; Davidowitz et al. 2004; Garrad et al. 2016). However, temperature is not the sole environmental variable that changes across latitude. Factors such as season length and food availability cannot be ruled out (Diamond and Kingsolver 2010; Horne et al. 2015). In other ectothermic species, season length has been found to have an equal or larger effect on body size than temperature (grasshoppers; Mousseau 1997; Walters and Hassall 2006). For treehole-breeding mosquitoes, season length is driven by rainfall, which, in the western U.S. tends to increase with latitude and therefore would drive the same patterns. For example, many areas in southern California receive less than 40 cm of rainfall annually during sporadic rain events in the winter and early spring, creating a very short season, whereas our most northern site in Oregon receives 120 cm of rainfall annually with a rainy season that extends into the summer. Mosquitoes at higher latitudes undergo larval diapause during winter, and this prolonged developmental period could produce larger individuals (Jordan and Bradshaw 1978).

Concordant with the body size results, we also found greater thermal tolerance in mosquitoes from lower latitudes and warmer climates (Figure 4, Figure 5). Based on previous literature indicating that acclimation to warm temperatures increases thermal tolerance (Hoffmann et al. 2003; Angilletta 2009; Gray 2013), we hypothesize that this variation in thermal tolerance also may be driven primarily by plasticity and less due to genetically-based differences. While we did not directly measure variation in thermal tolerance among our laboratory-reared populations, other studies testing for local adaptation in thermal performance of mosquitoes and *Drosophila* have found minimal or no significant differences (Couper et al. 2024, Klepsatel et al. 2013). In our study this direct effect of temperature, likely operating through acclimation to the environment during larval stages, appears to have a more substantial impact on thermal tolerance than the opposing effect of reduced adult body size.

Body size plays an important role in modulating an organism’s experienced temperature (Pinsky 2019). Often, a positive correlation is observed between body mass and thermal tolerance (Ribeiro et al. 2012; Baudier et al. 2015; Klockmann et al. 2017); however, this is not universal, as some studies have reported the opposite effect, where smaller size confers higher thermal tolerance (Peck et al. 2009; Franken 2017). While a common explanation is that larger organisms have greater thermal inertia, thereby enhancing their ability to withstand temperature extremes, this mechanism is less applicable to small-bodied ectotherms, which weigh only a fraction of a gram. Recognizing this, alternative explanations include larger individuals having lower mass-specific metabolic rates and greater energy reserves, enhancing their resilience to temperature extremes (Leiva et al. 2024). In our study, the SEM revealed positive effects of size at the individual and population level on knockdown time. This suggests that shrinking sizes as a consequence of development at warmer temperatures could be maladaptive, leading to reduced thermal tolerances (Figure 5). However, our specified SEM is one of the possible theoretical models, and future experiments that more clearly separate the causal influence of size are needed.

For both size and thermal tolerance, females showed a stronger response across temperature, as there was a significant interaction between sex and temperature. That is, female body size increased more rapidly with increasing latitude, leading to increased sexual size dimorphism with increasing size. As females are the larger sex in mosquitoes, this pattern does not follow Rensch’s rule, which states that the sexual size dimorphism should be more pronounced the smaller the size of the species. However, Rensch’s rule is typically applied to *interspecific* variation rather than *intraspecific* variation (but see Blanckenhorn et al. 2006) and our study was not designed to rigorously test this hypothesis (Meiri and Liang 2021). We also found that there was a significant interaction between sex and source temperature in which females from warm temperatures are especially thermally tolerant. A meta-analysis of ectotherms in the wild found that females have greater heat tolerance plasticity than males, and that this is associated with differences in body mass between sexes (Pottier et al. 2021).

The findings of this study contribute to our understanding of the complex effects of the environment on morphological and physiological traits in mosquitoes. We report that size varies by 2.3 mm (120% relative to the smallest individual) and thermal tolerance varies by 34 minutes (132% relative to the earliest knockdown time) across a species range spanning over 10° of latitude and that this variation in size appears to be plastic in response to environmental conditions, likely involving temperature, rather than fixed genetic differences. Such plasticity has important implications for the fecundity, survival, feeding behavior, and other traits of individuals, as well as for the sensitivity of entire populations to future climate warming. For mosquitoes, variation in thermal tolerance is likely to influence their geographic range and capacity to vector diseases (Sternberg and Thomas 2014; Mordecai et al. 2019). While we document higher thermal tolerance in warmer, lower latitude populations, it remains unclear whether this is enough to counteract the future impacts of climate change.

