## Supplement for "Plasticity in mosquito size and thermal tolerance across a latitudinal climate gradient"

Table S1. Results of the LMMs that include latitude as an explanatory variable for the size and knockdown time of adults collected as pupae and the size of adults reared their entire lives in controlled temperatures. Population was included as a random effect (sigma=0.23 for a, sigma=1.14 for b, sigma=0.05 for c). Block was included as a random effect (sigma=1.06 for b).

| Term | Estimate | SE | η^2^ | df | χ² | p-value |
| --- | --- | --- | --- | --- | --- | --- |
| (a) impact on size of field-collected individuals | | | | | | |
| latitude | 0.06 | 0.02 | 0.44 | 1 | 7.6 | 0.006 |
| sex | 0.5 | 0.21 | 0.45 | 1 | 250.9 | <0.0001 |
| latitude x sex | -0.02 | 0.01 | 0.04 | 1 | 19.0 | <0.0001 |
| (b) impact on knockdown time | | | | | | |
| latitude | -0.60 | 0.11 | 0.57 | 1 | 14.1 | 0.0002 |
| sex | -8.03 | 3.44 | 0.004 | 1 | 1.7 | 0.19 |
| latitude x sex | 0.20 | 0.09 | 0.01 | 1 | 4.8 | 0.03 |
| size | 0.90 | 0.72 | 0.007 | 1 | 1.7 | 0.19 |
| (c) impact on size of lab-reared individuals | | | | | | |
| latitude (collected) | 0.02 | 0.01 | 0.44 | 1 | 5.2 | 0.02 |
| sex | -0.92 | 0.42 | 0.61 | 1 | 354.5 | <0.0001 |
| latitude (collected) x sex | 0.01 | 0.01 | 0.002 | 1 | 1.0 | 0.31 |
| temperature (reared) | -0.13 | 0.01 | 0.36 | 1 | 166.8 | <0.0001 |

Table S2. Results of the linear models on the wing to thorax ratio of all individuals collected as pupae (N=423) and individuals in a single population reared their entire lives in controlled temperatures (N=28). Population was included as a random effect in (a) (sigma=0.03).

| Term | Estimate | SE | η^2^ | df | χ² | p-value |
| --- | --- | --- | --- | --- | --- | --- |
| (a) impact on wing to thorax ratio of field-collected individuals | | | | | | |
| temperature (collected) | -0.0007 | 0.01 | 0.001 | 1 | 0.01 | 0.91 |
| sex | 0.003 | 0.02 | 0.007 | 1 | 0.03 | 0.86 |
| (c) impact on wing to thorax ratio of lab-reared individuals | | | | | | |
| temperature (reared) | 0.0001 | 0.02 | 0.01 | 1 | 0 | 0.99 |
| sex | 0.02 | 0.04 | 0.001 | 1 | 0.3 | 0.56 |

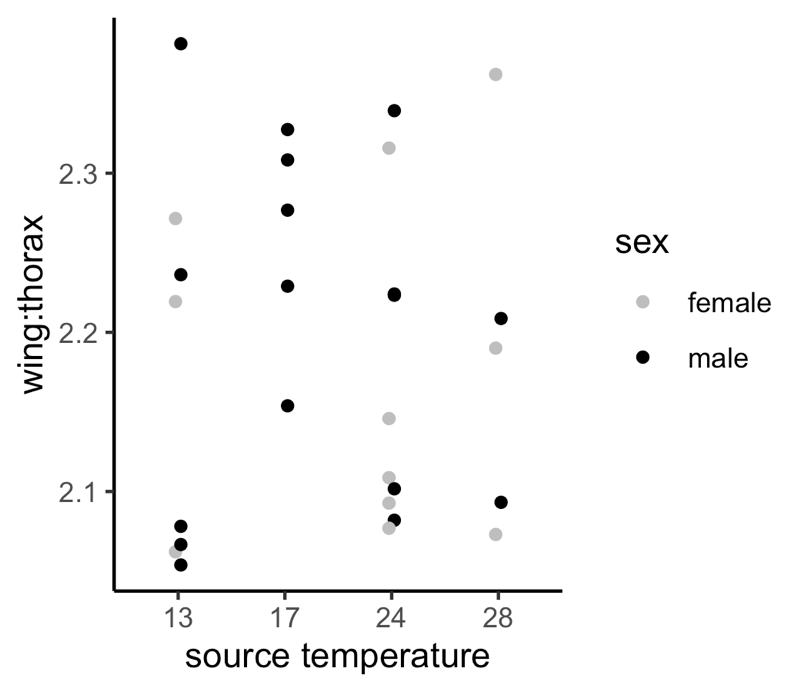

Figure S1. The temperature at which the mosquitoes were reared in the common garden experiment did not significantly impact the wing to thorax ratio.
